# Resolving widespread incomplete and uneven archaeal classifications based on a rank-normalized genome-based taxonomy

**DOI:** 10.1101/2020.03.01.972265

**Authors:** Christian Rinke, Maria Chuvochina, Aaron J. Mussig, Pierre-Alain Chaumeil, Adrian A. Davin, David W. Waite, William B Whitman, Donovan H. Parks, Philip Hugenholtz

## Abstract

An increasing wealth of genomic data from cultured and uncultured microorganisms provides the opportunity to develop a systematic taxonomy based on evolutionary relationships. Here we propose a standardized archaeal taxonomy, as part of the Genome Taxonomy Database (GTDB), derived from a 122 concatenated protein phylogeny that resolves polyphyletic groups and normalizes ranks based on relative evolutionary divergence (RED). The resulting archaeal taxonomy is stable under a range of phylogenetic variables, including marker genes, inference methods, corrections for rate heterogeneity and compositional bias, tree rooting scenarios, and expansion of the genome database. Rank normalization was shown to robustly correct for substitution rates varying up to 30-fold using simulated datasets. Taxonomic curation follows the rules of the International Code of Nomenclature of Prokaryotes (ICNP) while taking into account proposals to formally recognise the rank of phylum and to use genome sequences as type material. The taxonomy is based on 2,392 quality screened archaeal genomes, the great majority of which (93.3%) required one or more changes to their existing taxonomy, mostly as a result of incomplete classification. In total, 16 archaeal phyla are described, including reclassification of three major monophyletic units from the former Euryarchaeota and one phylum resulting from uniting the TACK superphylum into a single phylum. The taxonomy is publicly available at the GTDB website (https://gtdb.ecogenomic.org).

## Main

Carl Woese’s discovery of the Archaea in 1977, originally termed Archaebacteria (Woese and Fox, 1977) gave rise to the recognition of a new domain of life and fundamentally changed our view of cellular evolution on Earth. In the following decades, an increasing number of archaea were described, initially from extreme environments but subsequently also from soils, oceans, freshwater, and animal guts, highlighting the global importance of this domain (Gribaldo and Brochier-Armanet, 2006). Since their recognition, archaea have been classified primarily via genotype, i.e. small subunit (SSU) rRNA gene sequences, and hence, compared to bacteria, they suffer less from historical misclassifications based on phenotypic properties (Zuo et al., 2015). Using the SSU rRNA gene, Woese initially described two major lines of archaeal descent, the Euryarchaeota and the Crenarchaeota (Woese et al., 1990), and in the following years all newly discovered archaeal lineages were added to these two main groups. Non-extremophilic Archaea were generally classified as Euryarchaeota (Spang et al., 2017), which together with the discovery of novel extremophile euryarchaeotes (Baker et al., 2020) contributed to a considerable expansion of this lineage. Eventually two new archaeal phyla were proposed based on phylogenetic novelty of their SSU rRNA sequences; the Korarchaeota (Barns et al., 1996), recovered from hot springs in Yellowstone National Park, and the nanosized, symbiotic Nanoarchaeota (Huber et al., 2002), co-cultured from a submarine hot vent. By the mid-2000s, archaeal classification had begun to leverage genome sequences and the first sequenced crenarchaeote, *Ca*. Cenarchaeum symbiosum from a marine sponge (Hallam et al., 2006), was used to argue that mesophilic archaea should be considered as a separate phylum from hyperthermophilic Crenarchaeota for which the name Thaumarchaeota was eventually proposed (Brochier-Armanet et al., 2008). Subsequently the field experienced a burst in availability of genomic data due to the substantial acceleration of culture-independent genome recovery driven by improvements in high throughput sequencing (Adam et al., 2017; Spang et al., 2017). This resulted in the description and naming of several new archaeal lineages, including the candidate phyla Aigarchaeota (Nunoura et al., 2011), Geoarchaeota (Kozubal et al., 2013) and Bathyarchaeota (Meng et al., 2014), previously reported as members of the Crenarchaeota based on SSU rRNA data. Some of the proposed phyla were met with criticism, such as the Geoarchaeota, which was considered to be a member of the order Thermoproteales rather than a novel phylum (Guy et al., 2014). All former crenarchaeal lineages (Adam et al., 2017) were then recombined into the TACK superphylum, originally comprising the Thaumarchaeota, Aigarchaeota, the remaining Crenarchaeota, and Korarchaeota (Guy and Ettema, 2011), and more recently the Verstraetearchaeota (Vanwonterghem et al., 2016).

New archaeal lineages were also described outside the Euryarchaeota and TACK, which were ultimately classified into two superphyla, DPANN and Asgard (Rinke et al., 2013; Zaremba-Niedzwiedzka et al., 2017). DPANN was originally proposed based on five phyla; Diapherotrites, Parvarchaeota, Aenigmarchaeota, Nanoarchaeota and Nanohaloarchaeota (Rinke et al., 2013), but now also includes the Micrarchaeota, Woesearchaeota, Pacearchaeota, Altiarchaeota and Huberarchaeota (Adam et al., 2017; Baker et al., 2010; Castelle et al., 2015; Probst et al., 2018, 2014). The Asgard archaea are notable for their inferred sister relationship to the eukaryotes and were originally proposed based on the phyla Lokiarchaeota, Thorarchaeota, Odinarchaeota and Heimdallarchaeota (Seitz et al., 2016; Spang et al., 2015; Zaremba-Niedzwiedzka et al., 2017), followed by the Helarchaeota (Seitz et al., 2019). The net result of these cumulative activities is that archaeal classification at higher ranks is currently very uneven. The Euryarchaeota absorbed novel lineages and grew into a phylogenetic behemoth, whereas the Crenarchaeota were split into multiple subordinate taxa including several shallow lineages, such as Geoarchaeota, which despite their phylogenetic disproportion have been given the same rank of phylum. Attempts to rectify this taxonomic bias included a proposal to reclassify TACK as a single phylum termed Proteoarchaeota (Petitjean et al., 2014) and to introduce a new taxonomic rank above the class level which would generate several superclasses within the Euryarchaeota (Adam et al., 2017; Petitjean et al., 2015).

Most of this activity, i.e. the proposing and naming of new phyla, has occurred outside the International Code of Nomenclature of Prokaryotes (Parker et al., 2019) because the ICNP does not currently extend to uncultured microorganisms nor recognise the ranks of phylum and superphylum. However, proposals have been made to include the rank of phylum (Oren et al., 2015) and to allow gene sequences to serve as type material (Whitman, 2016), which have been increasingly adopted informally by the research community (Chuvochina et al., 2019). Currently, uncultured archaea and bacteria can be provisionally named using *Candidatus* status (Murray and Stackebrandt, 1995). However, these names have no formal standing in nomenclature, do not have priority, and often contradict the current ICNP rules or are otherwise problematic (Oren, 2017). The waters have been further muddied by the proposal of names for higher taxa without designation of lower ranks and type material, which leaves them without nomenclatural anchors and their circumscription subject to dispute if they become polyphyletic (Chuvochina et al., 2019). In addition to these higher level classification issues, the current archaeal taxonomy suffers from the same phylogenetic inconsistencies observed in the Bacteria, such as polyphyletic taxa (e.g. class Methanomicrobia; see below), but to a lesser degree than the Bacteria due to the early integration of phylogeny and the relatively small size of the archaeal dataset. More problematic is the widespread incomplete classification of environmental archaeal sequences in the NCBI taxonomy, which are often only assigned to a candidate phylum with no subordinate rank names. This high degree of incomplete classification is likely due to a natural hesitancy to create novel genera and intermediate taxa for groups lacking isolated representatives. Further, it is our opinion that the most useful biological classification should be based on an evolutionary framework, which takes into account differing rates of evolution such that ancestral taxa of a given rank co-existed in time (Parks et al., 2018). Currently, differences in evolutionary rates are not explicitly taken into account in the NCBI taxonomy.

For the domain Bacteria these long-standing taxonomic issues and inconsistencies were recently addressed by proposing a standardized taxonomy referred to as the Genome Taxonomy Database (GTDB; gtdb.ecogenomic.org). The GTDB normalizes rank assignments using relative evolutionary divergence (RED) in a genome phylogeny, which takes into account differing evolutionary rates, followed by an extensive automated and manual taxonomy curation process. This approach resulted in taxonomic changes to over half of the nearly 95,000 analyzed bacterial genomes (Parks et al., 2018) and has recently been extended to a complete classification from domain to species (Parks et al., 2019). Here we present the GTDB taxonomy for the domain Archaea (release R04-RS89), comprising 2,392 quality screened genomes from cultivated and uncultured organisms. This taxonomic release circumscribes 16 phyla, including three phyla from major monophyletic units of the Euryarchaeota, and one phylum resulting from the amalgamation of the TACK superphylum. The archaeal GTDB taxonomy is publicly available at the Genome Taxonomy Database website (https://gtdb.ecogenomic.org/).

## Results

### Reference genome tree and initial decoration with the NCBI taxonomy

The archaeal Genome Taxonomy Database (GTDB; 04-RS89) currently comprises 2392 quality-filtered genomes obtained from RefSeq/GenBank release 89 (*see Methods*, (Haft et al., 2018). Genomes were clustered into species units based on average nucleotide identity (ANI; *see Methods*), resulting in 1248 species (Parks et al., 2019). A representative sequence of each species, i.e. the genome of the type strain or the highest quality genome of an uncultivated species, was then used for phylogenomic analyses (**Table S1**). The protein sequences for up to 122 conserved single-copy ubiquitous archaeal genes were recovered from each genome (Parks et al., 2017); **Table S2, Fig. S1**), aligned, concatenated into a supermatrix, and trimmed to 5124 columns (*see Methods*). A reference tree (ar122.r89) was inferred using the C10 protein mixture model with the posterior mean site frequency (PMSF) approximation (Wang et al., 2018) implemented in IQ-TREE (Nguyen et al., 2015). The PMSF model is a faster approximation of finite mixture models, which capture the heterogeneity in the amino acid substitution process between sites, and is able to mitigate long-branch attraction artefacts (Wang et al., 2018). Additional trees were inferred from different alignments and with alternative inference methods to assess the robustness of the GTDB taxonomy (*see below*). The reference tree was initially decorated with taxon names obtained from the NCBI taxonomy (Federhen, 2012) standardized to seven canonical ranks as previously described (Parks et al., 2018). Strikingly, many archaeal genomes had no canonical rank information beyond their phylum affiliation (31.0%), which is partly offset by the extensive use of names with no rank in the NCBI taxonomy (**Fig. S2a**). This was particularly apparent for DPANN phyla, which almost entirely lack information in the family to class ranks (**Fig. S2b**).

### Removal of polyphyletic groups and rank normalization

Approximately 10% of NCBI-defined taxa above the rank of species (26 of 252) could not be reproducibly resolved as monophyletic or operationally monophyletic in the reference tree (the latter is defined as having an F measure ≥0.95; Parks et al., 2018; **Table S3;** *see Methods*). These include the phyla Nanoarchaeota, Aenigmarchaeota and Woesearchaeota, which were intermingled with each other and with unclassified archaeal genomes (**Fig. S3**). Another prominent example is the class Methanomicrobia, which comprised three orders in NCBI, two of which (Methanosarcinales and Methanocellales) were clearly separated from the type order Methanomicrobiales (**Fig. S4**.). To resolve these cases, the lineage containing the nomenclature type retained the name. Where possible, all other groups were renamed following the International Code of Nomenclature of Prokaryotes (ICNP) and recent proposals to modify the Code (*see Methods*). For example, the Methanomicrobia were resolved in GTDB by reserving the name for the lineage containing the type genus of the type order of the class, i.e. *Methanomicrobium*, and by reclassifying the two remaining orders into their own classes; Methanosarcinia class. nov. and Methanocellia class. nov. (**Fig. S4**). These and other ICNP-based rules used to standardize the curation process above the rank of genus are summarized in a decision tree (**Fig. S21, Suppl. Text**). When nomenclature types were not available at the genus and species level, the existing names were retained as placeholders with alphabetical suffixes indicating polyphyly. For example, the genus *Thermococcus* is polyphyletic because it also comprises species of the genus *Pyrococcus*. This polyphyly was resolved by retaining the name for the monophyletic group containing the type species, *Thermococcus celer*, and by assigning alphabetical suffixes to three basal groups comprising other *Thermococcus* genomes (**Fig. S5**). Note that *Thermococcus chitonophagus* was transferred to *Pyrococcus* according to the GTDB reclassification due to its proximity to the type species of this genus (**Fig. S5**).

Taxonomic ranks were normalized using the relative evolutionary divergence (RED) metric. This method linearly interpolates the inferred phylogenetic distances between the last common ancestor (set to RED=0) and all extant taxa (RED=1) providing an approximation of relative time of divergence (Parks et al., 2018). We tested the tolerance of the RED approach to increasing differences in substitution rates between lineages using simulated datasets. The approach was robust to variable rates up to 30-fold different (**Suppl. Text**), which is likely substantially higher than naturally occurring variation between prokaryotic taxa (Marin et al., 2017). Rank distributions in the NCBI-decorated reference tree were extremely broad, highlighting severely under- and over-classified outlier taxa (**Fig. 1a)**. Rank distributions were normalized (Parks et al., 2018) by systematically reclassifying outliers either by re-assignment to a new rank with associated nomenclatural changes for Latin names, or by moving names to new interior nodes in the tree (**Fig. 1b)**. This resulted in large movements of the median RED values of the higher ranks (order and above) to produce a normalized distribution (**Fig. 1b**). Over half (56.4%) of all archaeal NCBI taxon names had to be changed in the GTDB taxonomy, with the largest percentage of changes occurring at the phylum level (76.6%) (**Fig. 1c**). Examples of changes at lower ranks due to RED normalization include the genus *Methanobrevibacter*, which was divided into five genus-level groups; *Methanobrevibacter* which includes the type species and four genera with alphabetical suffixes (*Methanobrevibacter_A to Methanobrevibacter_D*; **Fig S6**). GTDB names were assigned to nodes with high bootstrap support (bs 98.5% ± 5.0%) to ensure taxonomic stability with a small number of exceptions (bs <90%) to preserve existing classifications (**Table S4**). Overall, 93.3% of the 2,239 archaeal genomes present in NCBI release 89 had one or more changes in their taxonomic assignments (**Fig. 1c**).

**Figure 1.**
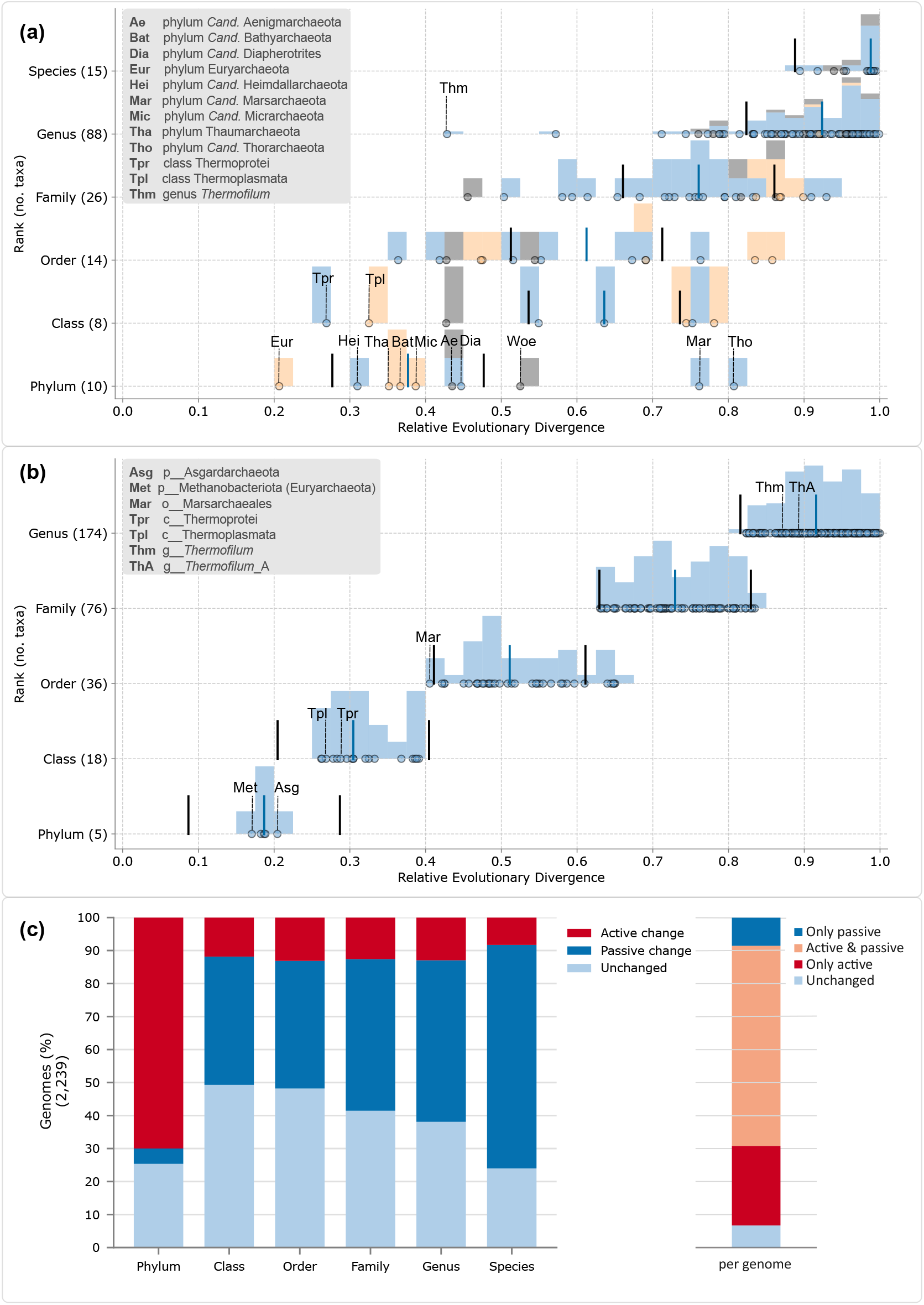
Comparison of rank normalized archaeal GTDB and NCBI taxonomies. **(a)** Relative evolutionary divergence (RED) of taxa defined by the NCBI taxonomy; **(b)** RED of taxa defined by the curated GTDB taxonomy (04-RS89). In (a) and (b) each data point (black circle) represents a taxon distributed according to its RED value (x-axis) and its rank (y-axis). The fill colours of the circle (blue, grey, or orange) indicate that a taxon is monophyletic, operationally monophyletic (defined as having an F measure >0.95), or polyphyletic, respectively in the underlying genome tree. An overlaid histogram shows the relative abundance of monophyletic, operationally monophyletic, and polyphyletic taxa for each 0.025 RED interval. A blue bar shows the median RED value, and two black bars the RED interval (+/- 0.1) for each rank. Note, that in the NCBI taxonomy the values of the higher ranks (order and above) are very unevenly distributed, to the point where the median distributions are not even in the expected order, i.e. the median RED value for classes was higher than the median order value. The GTDB taxonomy uses the RED value to resolve over- and under-classified taxa by moving them to a new interior node (horizontal shift in plot) or by assigning them to a new rank (vertical shift in plot). Only monophyletic or operationally monophyletic taxa were used to calculate the median RED values for each rank. In addition, only taxa with a minimum of two children (e.g. a phylum with two or more classes, or a class with two or more orders) were considered for the GTDB tree (--min_children 2), however, a more lenient approach (--min_children 0) was necessary for the NCBI tree since, except for the Euryarchaeota, none of the other NCBI phyla had the required minimum of two classes. Note, that the NCBI phylum Crenarchaeota is not displayed in the NCBI plot, since all genomes in this NCBI phylum are assigned to the class Thermoprotei, resulting in a single node decorated as “p Crenarchaeota; c Thermoprotei” (Tpr). Also, Korarchaeota are only represented by a single species, Korarchaeum cryptofilum, in r89 and hence there is no internal node to be displayed in this plot. RED values were calculated based on the ar122.r89 tree, inferred from 122 concatenated proteins, decorated with the NCBI and GTDB taxonomy, respectively. **(c)** Rank comparison of GTDB and NCBI taxonomies. Shown are changes in GTDB compared to the NCBI taxonomic assignments across 2,392 archaeal genomes from RefSeq/GenBank release 89. In the bars on the left, a taxon is shown as unchanged if its name was identical in both taxonomies, as a passive change if the GTDB taxonomy provided name information absent in the NCBI taxonomy (missing names), or as active change if the name was different between the two taxonomies. The right bar shows the changes of the entire tax string (consisting of seven ranks) per genome, indicating that most genomes had active and passive changes in their taxonomy. See Suppl. Text for more details.

### Robustness of proposed archaeal taxonomy

We tested the robustness of the monophyly and rank normalization of the proposed taxonomy in relation to a number of standard phylogenetic variables including marker genes, inference method, compositional bias, fast evolving sites, increasing number of genomes and rooting of the tree. Comparing tree similarities indicated that marker choice had a stronger influence on the tree topology than inference methods and substitution models (**Fig. S7; Fig. S8)**. However, for all subsequent comparisons we focused on the robustness of the taxonomy, not the overall consistency of tree topologies, as only a subset of interior nodes (69.9%) in the reference tree were used for taxon classification (**Table S5**).

#### Markers

As expected, individual protein phylogenies of the 122 markers had lower phylogenetic resolution than the reference tree, particularly at the higher ranks of class and phylum (**Fig S9a, b**). However, 78.5% of the GTDB taxa with ≥2 representatives above the rank of species were still recovered as monophyletic groups in ≥50% of single protein trees, and on average taxa were resolved as monophyletic in 74.1% of the single protein trees (**Fig 9c, d**). We also compared the GTDB taxonomy to SSU rRNA gene trees, due to their historical importance in defining archaeal taxa. However, this was complicated by the absence of this gene (>900 nt after quality-trimming) in almost half of the species representatives (578/1248; 46.3%). The majority (94.1%) of species representatives lacking SSU rRNA sequences were draft MAG assemblies (**Table S6**), which often lack this gene due to the difficulties of correctly assembling and binning rRNA repeats in metagenomic datasets (Parks et al., 2017; Sieber et al., 2018). Over 84% of the GTDB taxa with ≥2 SSU rRNA representatives were operationally monophyletic in the SSU rRNA tree, with loss of monophyly most pronounced in the higher ranks (**Fig. S10**) as seen for the single protein phylogenies. Three alternative concatenated protein marker datasets were assessed; rp1 (16 ribosomal proteins, **Table S7** (Hug et al., 2016), rp2 (23 ribosomal proteins, **Table S7**, (Rinke et al., 2013) and 53 recently proposed archaeal markers (ar.53; **Table S8** (Dombrowski et al., 2020). The great majority (≥95%) of GTDB taxa above the rank of genus with ≥2 representatives were recovered as monophyletic groups in the three alternative marker set trees inferred using the same C10 PMSF mixture model as the reference tree (**Fig. 4; Fig. S11**) (Rinke et al., 2013). The small percentage of taxa not resolved in the alternative trees (**Fig S11; Table S9**) were well supported in the reference tree (average bootstrap support 90.3±8.8% ;**Fig. S12**). However, these taxa were resolved as operationally monophyletic in a lower proportion of individual protein trees than other GTDB taxa (33.7±14.7% vs 77.1±18.3% respectively). The RED distributions of the GTDB taxa were comparable in the reference, rp1 and rp2 trees, whereas the SSU rRNA tree had a substantially broader distribution (**Fig. 2a**,**e, Fig. S13**), likely reflecting undersampling of the topology, lower resolution of single marker genes relative to concatenated marker sets and potentially compositional bias in the SSU rRNAs of thermophiles (Galtier & Lobry, 1997).

**Figure 2.**
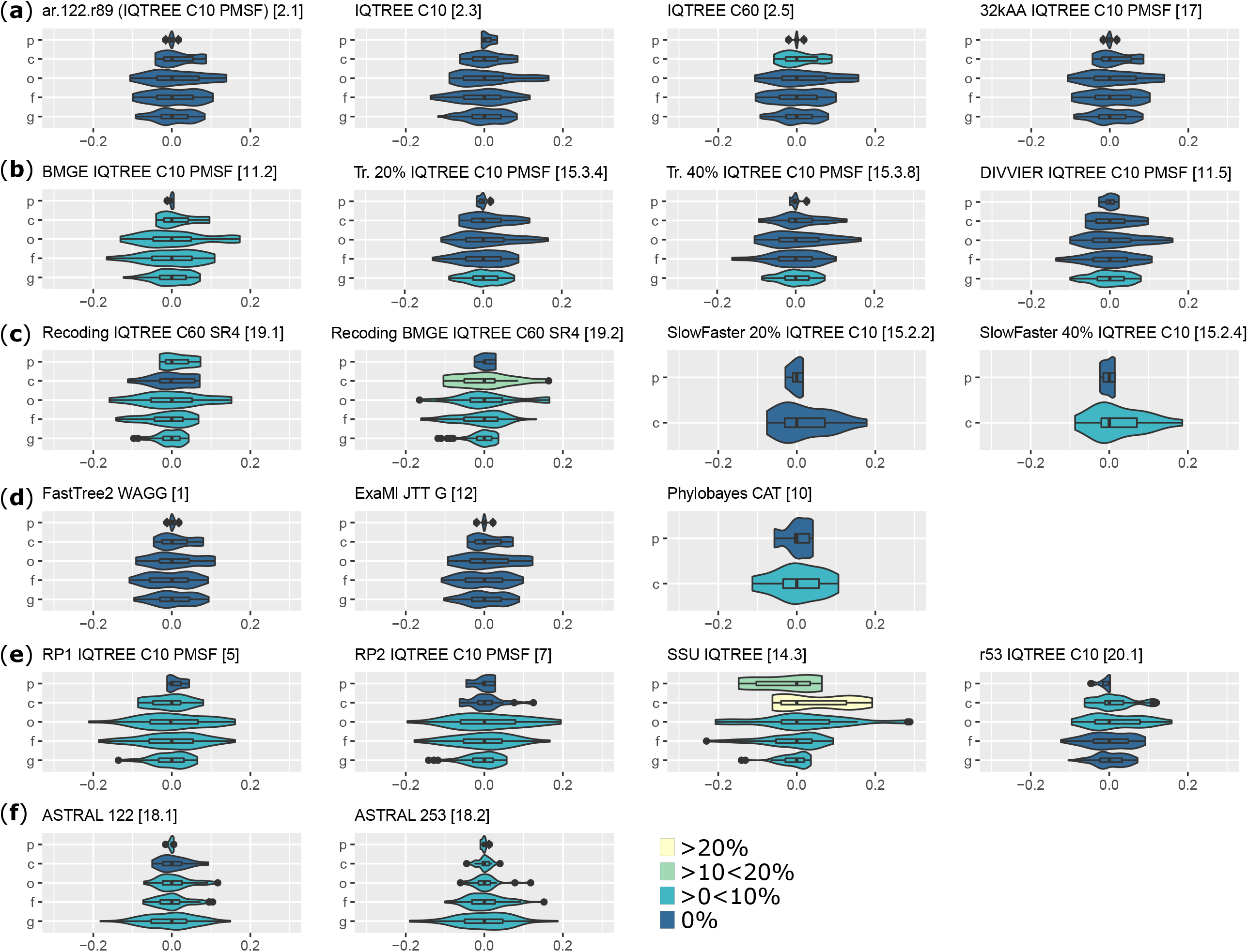
Comparison of marker sets, inference methods and models. Phylogenetic trees inferred with different methods, from varying concatenated alignments, or via supertree approaches were decorated with the GTDB 04-RS89 taxonomy. RED distributions for taxa at each rank (p, phylum; c, class; o, order; f, family; g, genus) are shown relative to the median RED value of the rank. The legend indicates the percentage of polyphyletic taxa per rank, defined as an F measure <0.95. Note that only taxa with two or more genomes were included. **(a)** trees inferred with IQ-TREE from a concatenated alignment of the 122 GTDB markers with ∼5,000 alignment columns using different profile mixture models (C10 PMSF, C10, C60), and from the untrimmed alignment 32kAA alignment. **(b)** Trees inferred from a modified concatenated alignment of the 122 GTDB markers to account for compositional bias, including stationary (BMGE) and progressive trimming (Tr. 20%, Tr. 40%) of heterogeneous sites, and clustering of sites with shared homology (Divvier). **(c)** Trees inferred from a modified concatenated alignment of the 122 GTDB markers including recoding into 4 character-states (C60 SR4), recoding and stationary trimming (BMGE C60 SR4), and removal of 20% and 40% of the fastest evolving sites (SlowFaster 20% and 40%). Note that, due to technical limitations a reduced order-dereplicated genome set was used for SlowFaster, allowing the evaluation of the ranks phylum and class only. **(d)** Trees inferred from 122 marker alignments using different inference software and models, including FastTree2, ExaML, and Phylobayes Note due to computational constrains PhyloBayes was calculated from the order dereplicated data set, allowing the evaluation of the ranks phylum and class only. **(e)**Trees inferred from alternative markers, including 16 ribosomal proteins (rp1), 23 ribosomal proteins (rp2), SSU rRNA genes (SSU), and a set of 53 marker proteins (ar53). **(f)** Trees inferred with the ASTRAL supertree approach using 122 and 253 marker proteins. More details about inference models and methods are given in **Table S10**. The number in square brackets following each tree name, e.g. [2.1] refers to the number of this tree in the supplements, including Table S10.

#### Inference methods and models

We overlaid the IQ-TREE-based taxonomy onto trees inferred with different models (e.g. C20, C60; **Table S10**) and phylogenetic inference tools, including FastTree and ExaML (maximum likelihood), PhyloBayes (Bayesian), and ASTRAL (supertree). All methods were applied to the 122 archaeal marker set with the exception of the ASTRAL supertree, which was also applied to a 253 concatenated marker set, subsampled from the PhyloPhlAn dataset (Segata et al., 2013). Overall, the GTDB taxonomy was remarkably consistent with comparable RED distributions for taxa at each investigated rank (**Fig. 2a**,**d; Fig. S13**). All GTDB taxa were recovered as monophyletic or operationally monophyletic (**Fig S11**) for all supermatrix methods, regardless of the underlying inference algorithm or model. The ASTRAL supertrees, inferred using the most divergent approach tested, recovered 96% of GTDB taxa with ≥2 representatives as monophyletic or operationally monophyletic (**Fig. S11**). The only major inconsistency affected the Euryarchaeota, with 48 taxa being placed outside of this phylum in both supertrees (**Fig S14e, Fig S15**).

#### Compositional bias and fast evolving sites

To test for possible compositional bias we employed tools for character trimming and for clustering of high confidence positional homology (*see Methods*) which have been shown to alleviate long branch attraction artifacts and to increase tree accuracy for alignments of distantly-related sequences (Ali et al., 2019; Criscuolo and Gribaldo, 2010). Decorated ML trees calculated from the trimmed or clustered alignments were in strong taxonomic agreement with the reference tree, showing comparable RED values and recovering >96.5% of GTDB taxa (with ≥2 representatives) as monophyletic or operationally monophyletic (**Fig. 2b, c; Fig. S11; Fig S13**). Less than 4% of genomes had conflicting taxonomic assignments at any rank above species (**Table S11**), with the largest difference being observed in the class Methanosarcinia (18 taxa) and the order Desulfurococcales (46 taxa) for the character trimmed alignment (**Fig S14d**). The clustered alignment resulted in fewer differences, with a total of seven conflicting taxa (**Fig S14d**).

#### Analysis of increasing genome database size

An important variable in producing a genome-based taxonomy is the changing number of genome sequences which will continue to increase in future and likely impact the underlying tree topology. To assess the robustness of the GTDB taxonomy to this variable, we recapitulated the expansion of the archaeal genome database since 2015 by subsampling the 1248 archaeal genomes according to their NCBI release date resulting in 362, 528, 1035 and 1183 taxa for 2015 to 2018, respectively. Comparison of the taxonomy derived from these antecedent trees to the reference tree revealed that >99% of all named taxa were recovered as monophyletic groups suggesting that addition of new branches to the reference tree does not destabilise the subset of robust interior nodes used for GTDB taxonomic assignments. This suggests that the GTDB taxonomy should be stable with increasing dataset size.

#### Rooting effects

Rank normalization is sensitive to the placement of the tree root, which defines the last common ancestor (set to RED=0) (Parks et al., 2020, 2018), and can therefore potentially influence the resulting taxonomy. Since the rooting of both the archaeal and bacterial domains remains contested, GTDB uses an operational approach whereby the median of multiple plausible rootings is considered (Parks et al., 2018). We assessed the effect of fixed rooting of the archaeal domain on the taxonomy by testing two recently proposed archaeal root placements; the first, within the Euryarchaeota (as currently defined by NCBI) (Raymann et al., 2015) and the second, between the DPANN superphylum and all other Archaea (Williams et al., 2017). Overall, RED values were stable across the tested rooting scenarios and the intervals defining taxonomic ranks in GTDB were largely preserved (**Fig. 3**). As expected, a fixed root caused the taxa within the rooted lineages to be drawn closer to the root, although it is important to note that absolute RED values should not be compared between trees, or even between different rootings of the same tree. Rather the distribution of labelled nodes within a rank is the key metric, and by this metric all taxa in the rooted lineages were within their expected RED assigned rank intervals (**Fig S16; Fig. 3**).

**Figure 3.**
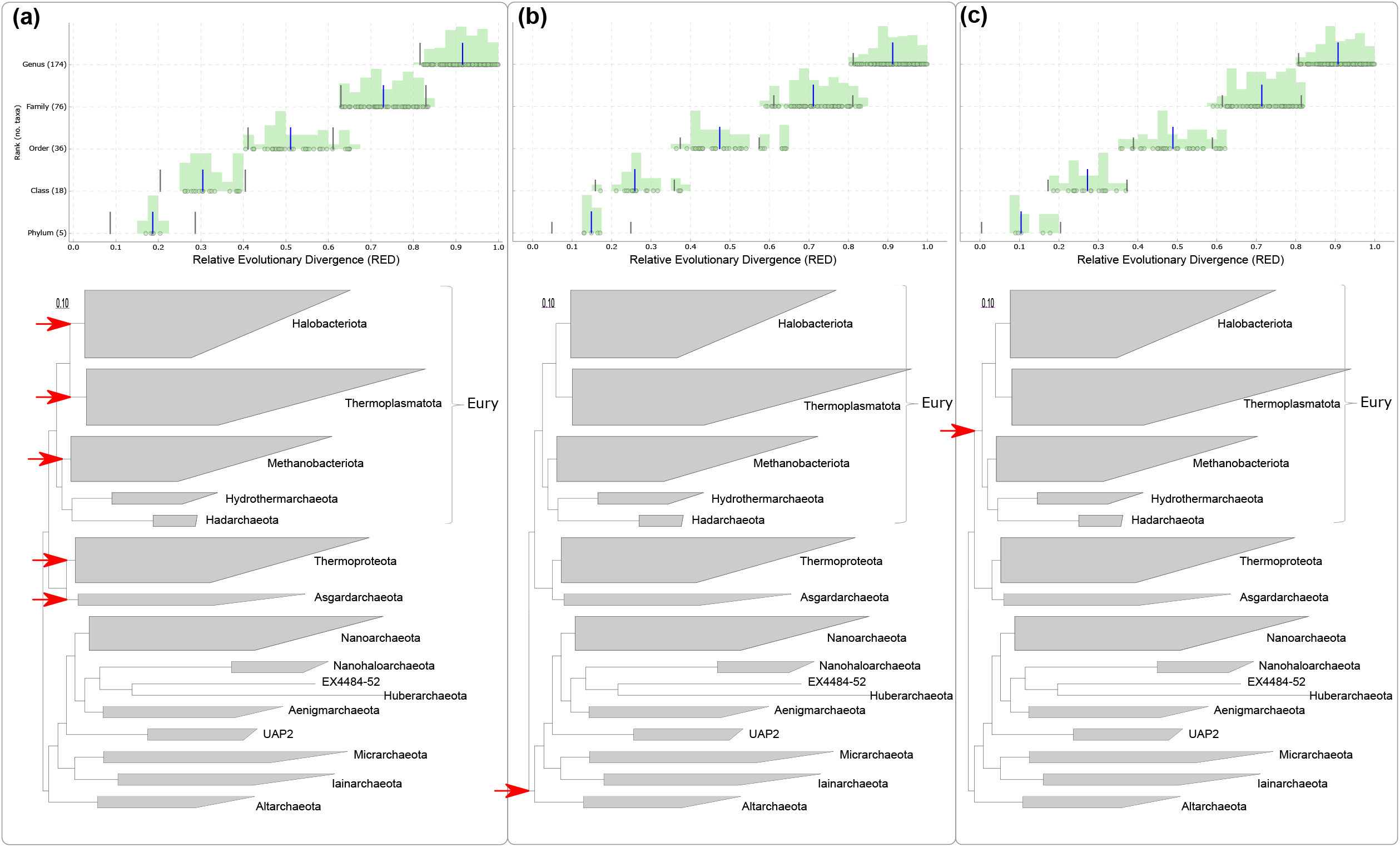
Impact of different rooting scenarios on the relative evolutionary divergence (RED). The rooting approach implemented in GTDB **(a)**, which calculates the relative evolutionary divergence (RED) as the median of all possible rootings of phyla with at least two classes (red arrows), is compared to a fixed root between the DPANN superphylum (red arrow) and the remaining Archaea **(b)**, and to a fixed root within the NCBI phylum Euryarchatoa, which translates to a root between the two phyla Thermoplasmata and Halobacteriota (red arrow) and the rest of the Archaea in the GTDB taxonomy **(c)**. In the upper RED plot each data point (black circle) represents a taxon distributed according to its RED value (x-axis) and its rank (y-axis). An overlaid histogram shows the relative abundance of taxa for each 0.025 RED interval, a blue bar shows the median RED value, and two black bars the RED interval (+/- 0.1) for each rank. Note that, overall, the ranks can be distinguished based on their RED value, regardless of the applied rooting scenario. Furthermore, RED values are relative and should not be directly compared between plots as they are dataset specific. Rather, the distribution of RED values, i.e. the distance (positive or negative) from the median of the RED value for each rank (ΔRED) is the key metric. The trees include a label highlighting the corresponding NCBI phylum Euryarchaeota (Eury), as a point of reference. The scale bars indicate 0.1 substitutions.

### Proposal of new and revised taxa based on the GTDB taxonomy

After resolving polyphyletic groups and normalizing ranks, the archaeal GTDB taxonomy (release R04-RS89) comprises 16 phyla, 36 classes, 96 orders, 238 families, 534 genera and 1248 species (**Table S12**). This entailed the proposal of 13 new taxa and 32 new candidatus taxa above the rank of genus including five novel species combinations and three novel candidatus species combinations (**Tables S13 to S16**). We also used 25 Latin names without standing in nomenclature as placeholders in the GTDB taxonomy (**Table S17)** to preserve literature continuity. The extensive rearrangement of phyla to normalize phylogenetic depth resulted in both division and amalgamation of release 89 NCBI phyla (**Fig. 4**). For example, the phylum Euryarchaeota was divided into five separate phyla in the GTDB taxonomy due to its anomalous depth (**Fig. 1a; Fig 4**). Names for three of the five phyla have previously been proposed; Methanobacteriota (Whitman et al., 2018), Hadarchaeota (Chuvochina et al., 2019), and Hydrothermarchaeota (Chuvochina et al., 2019). Names for the other two phyla are formally proposed in this study based on valid class names within each lineage; Halobacteriota phyl. nov. (after the class Halobacteria) and Thermoplasmatota *phyl. nov*. (after the class Thermoplasmata; **Table 1, Table S13**). Note that the name Euryarchaeota was not retained in the GTDB taxonomy for two reasons: Firstly the name would be illegitimate if the rank of phylum is introduced into the Code as a type was not designated in the original proposal (Woese et al., 1990), which is one of the requirements for validation of a name. Secondly, the substantial changes made to this phylum may introduce confusion if the name had been retained for one of the five newly circumscribed phyla. It is possible that the name Euryarchaeota could be reintroduced as a superphylum, however, this is outside of the GTDB taxonomy which only uses canonical ranks.

**Figure 4.**
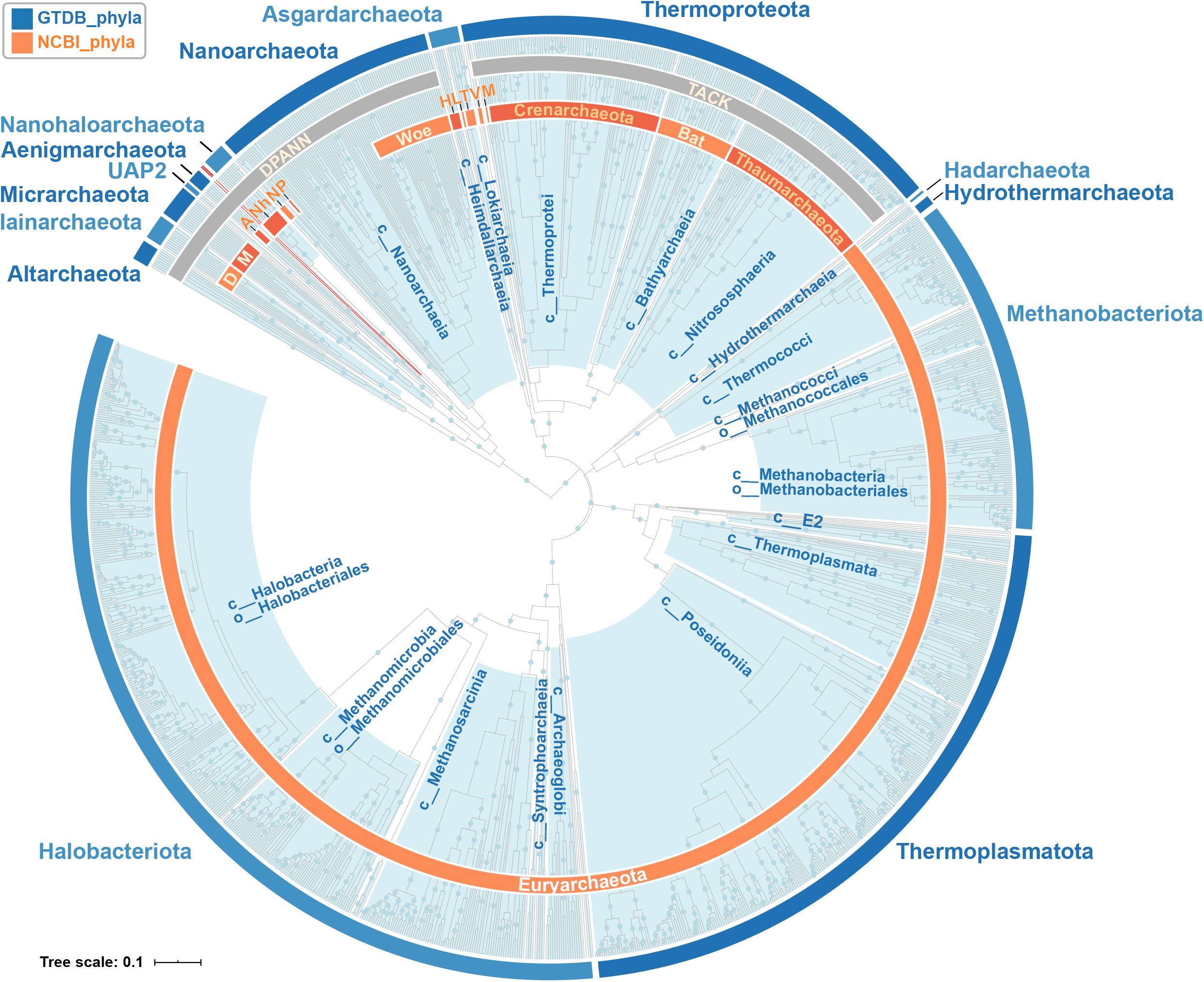
Rank normalized archaeal GTDB taxonomy. Species dereplicated scaled ar122.r89 tree decorated with the archaeal GTDB taxonomy R04-RS89. The outer blue ring denotes the rank normalized phyla, and the light blue clades indicate the classes in the rank normalized GTDB taxonomy. Classes with 10 or more taxa are labelled, and a below class divergence is indicated by providing the lower rank (e.g. order) for a given class. The two GTDB phyla consisting of only a single species each, namely Huberarchaeota and EX4484-52 are highlighted by red branches indicating their uncertain placement in the ar122.r89 tree. The inner orange ring denotes the r89 NCBI phyla with 2 or more taxa. The NCBI superphyla TACK and DPANN are indicated with gray color strips. Abbreviations are the following: Bat (*Ca*. Bathyarchaeota), M (*Ca*. Marsarchaeota), V (*Ca*. Verstraetearchaeota), T (*Ca*. Thorarchaeota), L (*Ca*. Lokiarchaeota), H (*Ca*. Heimdallarchaeota), Woe (*Ca*. Woesearchaeota), P (Ca. Parvarchaeota), N (Nanoarchaeota), Nh (*Ca*. Nanohaloarchaeota), A (*Ca*. Aenigmarchaeota), M (*Ca*. Micrarchaeota), D (*Ca*. Diapherotrites). Note: the scaled tree was generated by replacing the branch lengths with the median relative evolutionary distance (RED), calculated across all plausible rootings. Bootstrap values over 90% are indicated by blue dots. Scale bar indicates 0.1 RED.

**Table 1.**
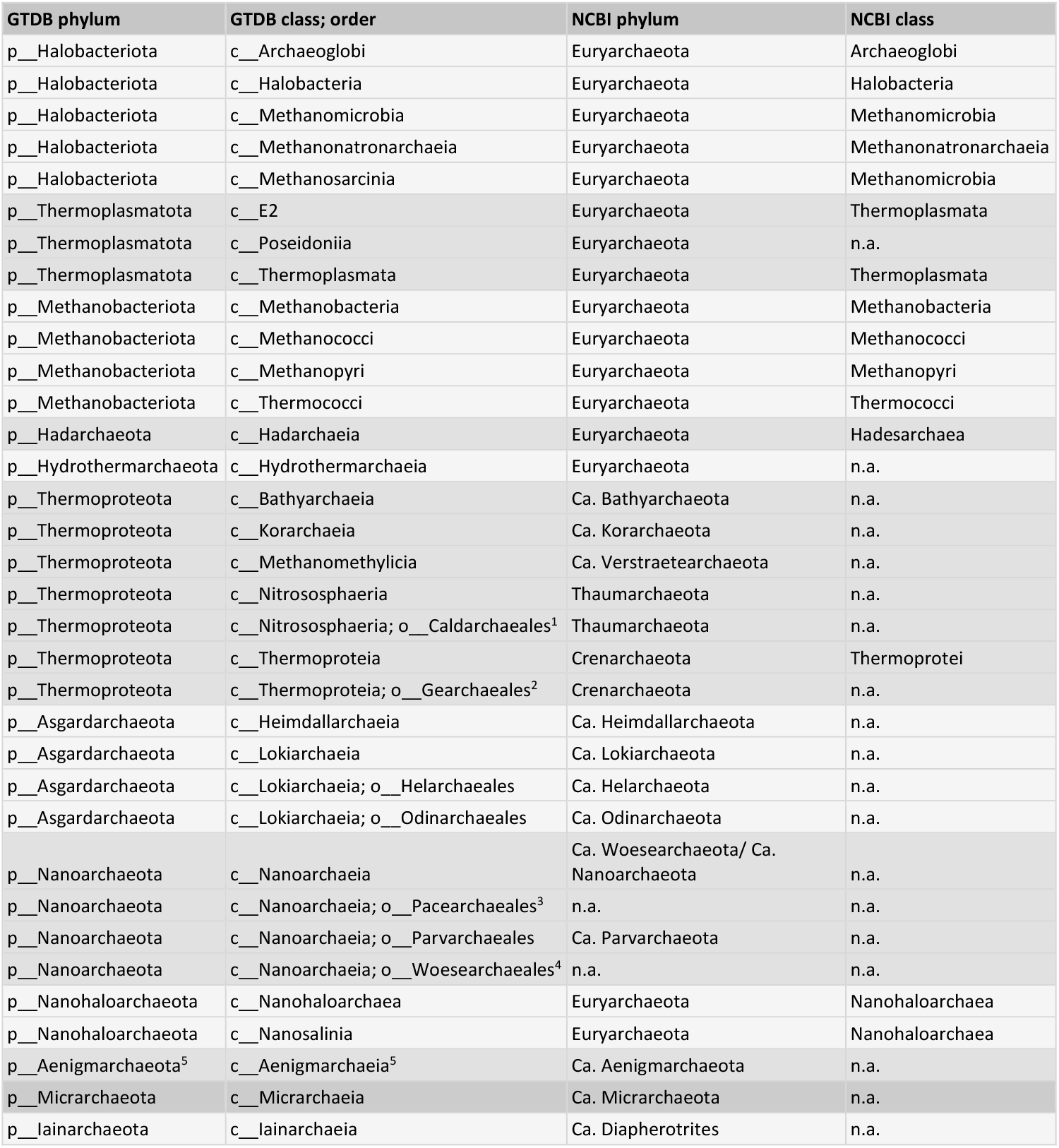
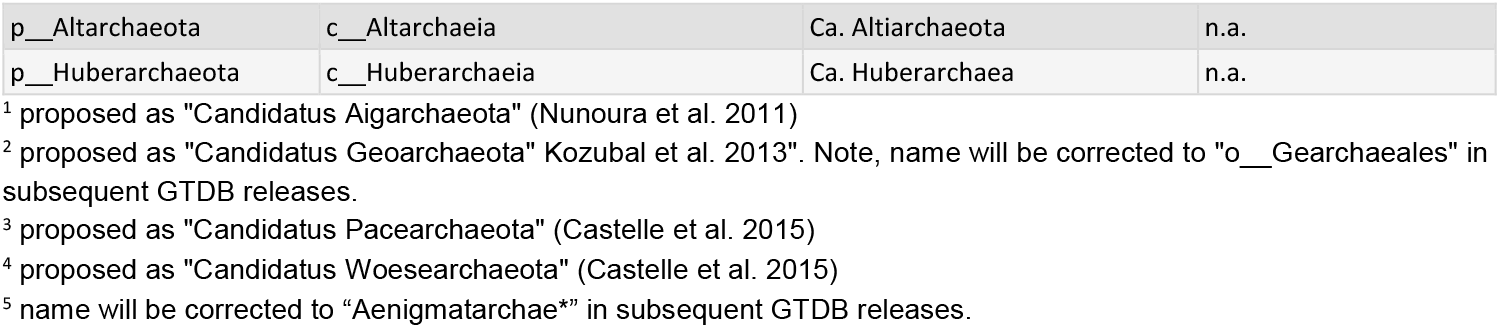
Correspondence between GTDB and NCBI taxonomy. Named GTDB phyla, major classes and selected orders are listed with their corresponding NCBI taxonomy. Note that, in cases where GTDB and NCBI lineages are not an exact match, the NCBI lineage with the highest number of matching taxa is provided. Abbreviations: n.a. = not assigned, meaning no rank has been assigned to this lineage in the NCBI taxonomy. Further details regarding the correspondence between NCBI and GTDB taxa are provided in Table S23 and S24.

The TACK superphylum (Guy and Ettema, 2011) was reclassified as a single phylum based on rank normalization and its robust monophyly, for which we propose to use the effectively published name Thermoproteota (Whitman et al., 2018) based on the earliest validly described class for any TACK phylum, the Thermoprotei (Reysenbach, 2001) (**Table 1, Table S13**). Note that alternatively the phylum name could have been derived from the earliest genus name in the TACK superphylum, *Sulfolobus*, however this genus is currently a member of the class Thermoprotei. The RED-based reorganisation of this lineage requires the unification of the NCBI-defined TACK phyla as previously proposed (Petitjean et al., 2014). Thus the Thaumarchaeota was reclassified as a class-level lineage, for which we propose an emended description of the only validly described class in this lineage, the Nitrososphaeria (Stieglmeier et al., 2014) (**Fig. 5)**. According to its RED-based rank and concatenated protein phylogeny, the Aigarchaeota constitutes an order-level lineage within the Nitrososphaeria, for which we propose the order Caldarchaeales *ord. nov*. (**Table S15**) derived from the species *Ca*. Caldarchaeum subterraneum (Nunoura et al., 2011). We propose to reclassify the NCBI-defined Crenarchaeota as an emended description of the class Thermoprotei and the Korarchaeota as the Korarchaeia *class. nov*. (**Table S15**) based on the type species *Ca*. Korarchaeum cryptofilum (Elkins et al., 2008). In addition, the candidate phyla Verstraetearchaeota (Vanwonterghem et al., 2016) and Bathyarchaeota (Meng et al., 2014) were unified with the Thermoproteota based on their robust phylogenetic affiliation. After unification, both lineages represent classes in the Thermoproteota for which the names *Ca*. Methanomethylicia (Oren et al., 2020) from *Ca*. Methanomethylicus mesodigestum (Vanwonterghem et al., 2016) and the GTDB placeholder name Bathyarchaeia (**Table S17**) were implemented, until type material is assigned for the latter lineage (Chuvochina et al., 2019). Note that the name Crenarchaeota was not retained in the GTDB taxonomy, *i*.*e*. used instead of the name Thermoproteota, to avoid confusion over its recircumscription relative to the NCBI taxonomy, and because the name would be illegitimate if the rank of phylum is introduced into the Code as per the Euryarchaeota (Woese et al., 1990).

**Figure 5.**
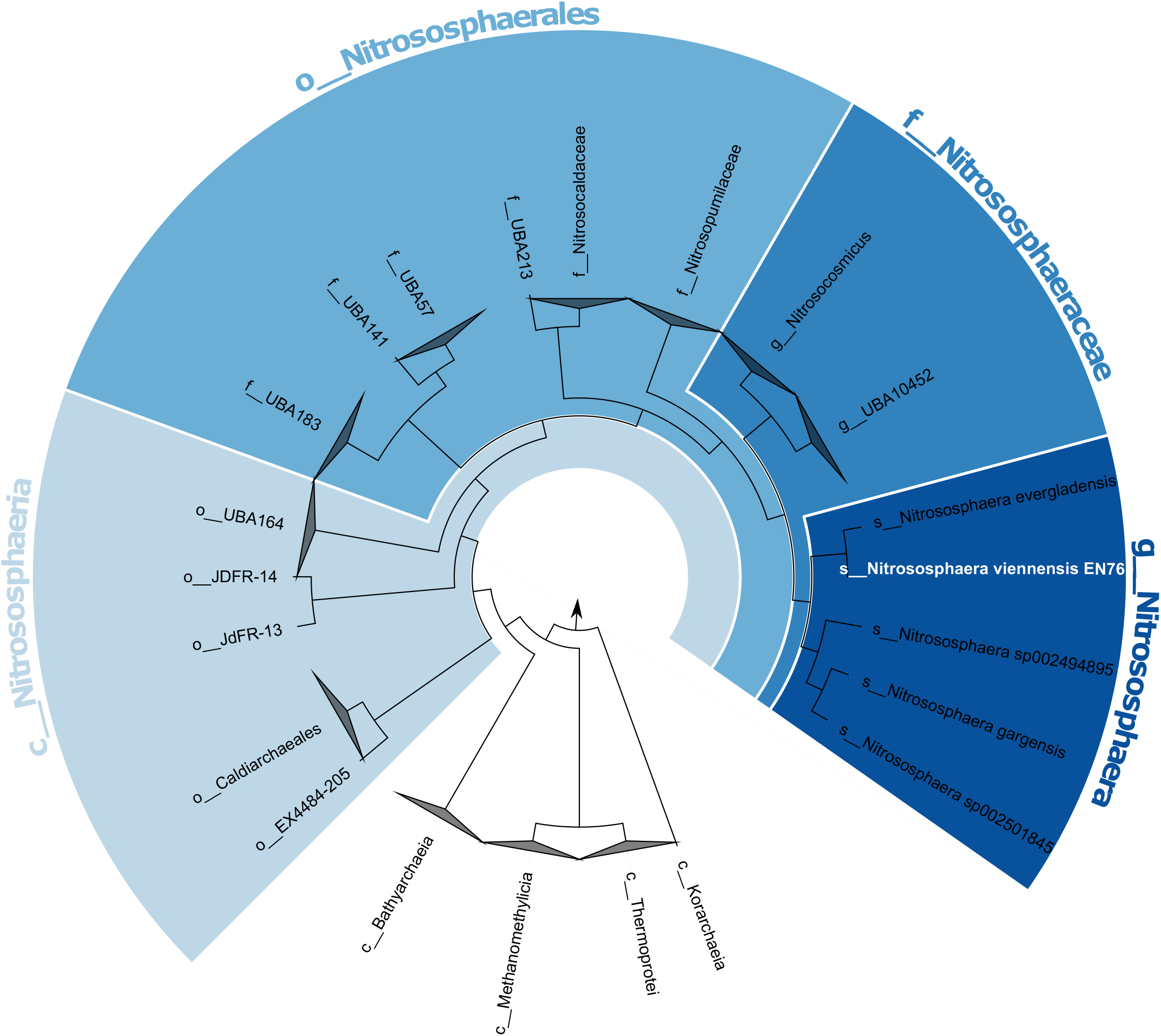
Reclassification of the members of the Thaumarchaeota. Cladogram based on the ar122.r89 reference tree showing the GTDB phylum Thermoproteota; with the classes Korarchaeia, Thermoprotei, Methanomethylicia (former phylum *Candidatus* Verstraetearchaeota), Bathyarchaeia, and Nitrososphaeria. The validly published class Nitrososphaeria (light blue) was emended in GTDB to include all taxa assigned to the phylum Thaumarchaeota in the NCBI r89 taxonomy. The type species of this lineage is *Nitrososphaera viennensis* (Stieglmeier et al. 2014), which serves as the type for higher taxa including the genus *Nitrososphaera*, the family *Nitrososphaeraceae*, the order *Nitrososphaerales*, and the class *Nitrososphaeria*. The genome of the *Nitrososphaera viennensis* type strain EN76^T^ (=DSM 26422^T^=JMC 19564^T^), is highlighted in white. Arrow points to outgroup.

Similar to the TACK superphylum, the recently described Asgard superphylum (Zaremba-Niedzwiedzka et al., 2017) was reclassified as a single phylum for which we use Asgardarchaeota as a placeholder name (**Table S17**) to retain its identity until type material has been proposed for this lineage (Chuvochina et al., 2019), which potentially could be the recently described co-culture, *Ca*. Prometheoarchaeon syntrophicum (Imachi et al., 2020). By contrast, the DPANN superphylum (Rinke et al., 2013) still comprises multiple (nine) phyla after rank normalization in part due to a lack of stable deeper interior nodes for naming (**Fig. 4**). Many of these phyla are represented by only a few genomes on long branches often reflected by low bootstrap support (**Fig. S17**). We envision that the DPANN phyla will undergo further reclassification in future GTDB releases when additional genomes become available to populate this region of the tree, however, even if some these phyla were unified, DPANN cannot be collapsed into a single phylum according to rank normalization.

Unlike the major changes required at the phylum level, archaeal taxa with lower rank information (species to class) were more stable, with an average of only 11% name changes. Notable examples include the well-known genus *Sulfolobus*, which was reported early on as potentially polyphyletic (Fuchs et al., 1996) and is comprised of strains differing in their metabolic repertoire and genome size (Quehenberger et al., 2017). The concatenated protein tree confirms that *Sulfolobus* is not monophyletic, being interspersed with species belonging to the genera *Acidanus, Metallosphaera* and *Sulfurisphaera*. We resolved this polyphyly by dividing the group into four separate genera (**Fig. S18**) reflecting previously reported differences between *Sulfolobus* species, including a high number of transposable elements in *Sulfolobus_A* species (*S. islandicus* and *S. solfataricus*), which are mostly absent in the type species of the genus, *Sulfolobus acidocaldarius* (Quehenberger et al., 2017). An example of a more complicated situation is the taxonomy of genera belonging to the halophilic family Natrialbaceae. Some of these genera have been reported as polyphyletic such as *Natrinema* and *Haloterrigena* (Minegishi et al., 2010), or are polyphyletic in published trees without associated comment, including *Halopiger* (Sorokin et al., 2019) and *Natronolimnobius* (Sorokin et al., 2018). We confirmed this polyphyly in the concatenated protein tree, which required extensive reclassification guided by the type species of each genus (**Fig S19, Table S14)**.

## Discussion

Here we present the Genome Taxonomy Database (GTDB) for the domain Archaea to provide a phylogenetically congruent and rank-normalized classification based on well-supported nodes in a phylogenomic tree of 122 concatenated conserved single copy marker proteins (**Table S2**). Unlike the bacterial GTDB reference tree comprising ∼19 fold more species representatives (Parks et al., 2020), the relatively modest number of publicly available archaeal genomes representing only 1248 species allowed us to use IQ-TREE with a protein mixture model that captures substitution site heterogeneity between sites and can mitigate long branch attraction artifacts (Wang et al., 2018). The proposed taxonomy is stable under a range of standard phylogenetic variables, including alternate marker genes, inference methods, tree rooting scenarios and expansion of the genome database, due in part to the inherent flexibility of rank designations within RED intervals (**Fig. 1**). We endeavoured to preserve the existing archaeal taxonomy wherever possible, however the great majority (93.3%) of the 2239 archaeal genomes in GTDB had one or more changes to their classification compared to the NCBI taxonomy (**Fig. 1c**). This high percentage of modifications, compared to 58% reported for genomes in the bacterial GTDB (Parks et al., 2018), could be attributed to extreme unevenness observed within archaeal ranks, particularly at the phylum level. For example, the division of the NCBI-defined Euryarchaeota alone affected over two thirds of all archaeal genomes (**Fig. 4**). Secondly, widespread missing rank information in NCBI r89, particularly amongst as-yet-uncultivated lineages, required numerous passive (rank filling) changes to the taxonomy (**Fig. 1**). And thirdly, the Bacteria have highly sampled species and genera that did not require taxonomic changes, including over 12,000 *Streptococcus* genomes in release 04-RS89, which effectively lowers the percentage of bacterial genomes with taxonomic differences to NCBI.

The 7-fold difference in the number of archaeal and bacterial phyla (16 vs 112; release 04-RS89) is notable and raises the question of whether GTDB-defined bacterial and archaeal phyla are comparable. Strictly speaking they are not, as each taxonomy was developed independently and shaped around the existing NCBI taxonomy as far as possible. Early attempts to produce a prokaryotic concatenated protein tree upon which to base a combined taxonomy distorted RED values to the point of being unusable (data not shown). In the case of the Archaea, we also experimented with moving the phylum and class rank intervals to the right (away from the root) since the starting distributions were so broad (Fig. 1a), which more than doubled the number of archaeal phyla. However, this also resulted in an even greater departure from the starting NCBI taxonomy and hence was not pursued further. Perhaps the most compelling evidence that the current ratio of phyla may reflect true biological diversity is the fact that there are ∼19-fold more ANI-defined bacterial than archaeal species (1248 vs 23,458; release 04-RS89).

The archaeal GTDB taxonomy and associated tree and alignment files are available online (https://gtdb.ecogenomic.org) and via linked third party tools, e.g. AnnoTree (Mendler et al., 2019). Users can classify their own genomes against the archaeal GTDB taxonomy using GTDB-Tk (Chaumeil et al., 2019). We envisage that in the short term, the archaeal taxonomy will scale with genome deposition in the public repositories based on current accumulation rates (**Fig. S20**). However, in the longer term more efficient phylogenomic tools will be needed to scale with increasingly large genomic datasets and to allow for biologically meaningful phylogenetic inferences. We also expect that less stable parts of the concatenated protein tree, *e*.*g*. the DPANN phyla, will become more robust with additional genome sequence representatives.

## Methods

### Genome dataset

For the archaeal GTDB taxonomy R04-RS89 we obtained 2,661 archaeal genomes from RefSeq/GenBank release 89 and augmented them with 187 phylogenetically diverse MAGs derived from the Sequencing Read Archive (SRA, (Haft et al., 2018)) as part of a large genome recovery study (Parks et al., 2017), resulting in 2,848 genomes. This data set was refined by applying a quality threshold (completeness - 5x contamination >50%) using lineage specific markers implemented in CheckM (Parks et al., 2019, 2015), and by screening out genomes which contain <40% of the 122 archaeal GTDB marker genes, more than 100,000 ambiguous bases, more than 1000 contigs, and which have an N50 <5kb. The filtered genomes were manually inspected and four exceptions were made for genomes of high nomenclatural or taxonomic significance (**Table S22)**. This approach left 2,392 genomes to form species clusters (see below), resulting in a total 1248 species representative genomes for the downstream analysis (**Table S1**). The 456 genomes which did not pass QC are still searchable on the GTDB website (https://gtdb.ecogenomic.org/) and are listed in **table S20**.

### NCBI taxonomy

The NCBI taxonomy of all representative genomes of R04-RS89 was obtained from the NCBI Taxonomy FTP site on July 16th, 2018. The NCBI taxonomy was standardized to seven ranks (species to domain) by identifying missing standard ranks and filling these gaps with rank prefixes and by removing non-standard ranks (Parks et al., 2018). All standard ranks were prefixed with rank identifiers (e.g. “p” for phylum) as previously described (McDonald et al., 2012).

### Phylogenomic marker set

Archaeal multiple sequence alignments (MSAs) were created through the concatenation of 122 phylogenetically informative markers comprised of proteins or protein domains specified in the Pfam v27 or TIGRFAMs v15.0 databases. The 122 archaeal marker proteins were selected based on the criteria described in (Parks et al., 2017). In brief, this included being present in ≥90% of archaeal genomes, and, when present, single-copy in ≥95% of genomes. Only genomes comprising ≤200 contigs with an N50 of ≥20 kb and with CheckM completeness and contamination estimates of ≥95% and ≤5%, respectively, were considered. Phylogenetically informative proteins were determined by filtering ubiquitous proteins whose gene trees had poor congruence with a set of subsampled concatenated genome trees (Parks et al., 2017). Gene calling was performed with Prodigal v2.6.3, and markers were identified and aligned using HMMER v3.1b1. The presence and absence of the 122 protein markers in the 1248 species representatives is provided in **table S21**. The marker proteins were concatenated into a multiple sequence alignment (MSA) of 32.5k columns, to which we refer as “untrimmed MSA” in this manuscript. To remove sites with weak phylogenetic signals, we created an amino acid alignment by trimming columns represented in <50% of the genomes and columns with less than 25% or more than 95% amino acid consensus, resulting in an initial 27,000 amino acid alignment (Suppl text). Thereby, the term consensus refers to the number of taxa with the same residue in a given sequence alignment column, e.g. a maximum consensus of 95% means that a maximum of 95% of all taxa can have the same residue in an alignment column in order to be considered for the trimmed alignment. To reduce computational requirements, the alignment was further trimmed by randomly selecting 42 amino acids from the remaining columns of each marker. The resulting ar122.r89 MSA included a total of 5124 (42*122) columns. This MSA filtering methodology is implemented in the `align` method of GTDB-Tk v1.0.2 (Chaumeil et al., 2019).

### Alternative supermatrix marker sets

Alternative MSAs, resembling previously published datasets, were created through the concatenation of 16 ribosomal proteins, termed dataset rp1 (Hug et al., 2016) and 23 ribosomal proteins, termed rp2 (Rinke et al., 2013). After trimming columns represented by <50% of the genomes and with an amino acid consensus <25%, the resulting MSA spanned 1174 and 2377 amino acids for rp1 and rp2, respectively.

In addition we created a MSA from 53 recently proposed archaeal marker proteins (ar.53; Table S8) (Dombrowski et al., 2020), by creating individual alignments for each of the 53 HMMs for all 1248 species representatives, using pfam 33.1 and tigrfam 15.0. The resulting concatenated MSA of 13,451 amino acids was used without further trimming steps to infer phylogenetic trees or to apply tools addressing compositional bias.

### SSU rRNA gene

Archaeal SSU rRNA genes were identified from the 1248 archaeal R04-RS89 GTDB genomes using nhmmer v3.1b2 (Wheeler and Eddy, 2013) with the SSU rRNA model (RF00177) from the RFAM database (Kalvari et al., 2018). Only the longest sequence was retained per genome. The resulting sequences were aligned with SSU-ALIGN 0.1.1 (Nawrocki, 2009), regions of low posterior probabilities, which are indicative of high alignment ambiguity were pruned with ssu-mask (SSU-ALIGN 0.1.1). The alignment was trimmed with an in-house script (trimSeqs.py v0.0.1; module bioscripts) to remove poorly represented leading and trailing positions, along with short sequences below 900bp (https://github.com/Ecogenomics/scripts/blob/master/scripts/trimSeqs.py).

### Species cluster

The 2,392 archaeal genomes were formed into species clusters as previously described (Parks et al., 2020). Briefly, a representative genome was selected for each of the 380 validly or effectively published archaeal species with one or more genomes passing quality control and genomes assigned to these representatives using average nucleotide identity (ANI) and alignment fraction (AF) criteria. Thereby, genomes were assigned to the closest representative genome for which they have an ANI of 95% ANI, with an AF of at least 65%, except if two representatives had an ANI >95%. In such cases, the ANI radius of a representative was set to the value of the closest representative up to a maximum of 97%, with species representatives having an ANI >97% considered synonyms. Genomes not assigned to one of these 380 species were formed into 868 *de novo* species clusters, each specified by a single representative genome. Representative genomes were selected based on assembly quality with preference given to isolate genomes. Of the 868 de novo species clusters, the representative genomes of 70 were unnamed isolates, 775 were MAGs and 23 were SAGs.

### Accounting for compositional bias

Each of the untrimmed 122 GTDB r89 archaeal single-copy marker protein alignments was filtered individually using BMGE 1.12 (Criscuolo and Gribaldo, 2010) and Divvier 1.0 (Ali et al., 2019). BMGE was executed using -t AA -s FAST -h 0.55 -m BLOSUM30, and Divvier was run using the recommended options: -divvy -mincol 4 -divvygap. Processing untrimmed protein alignments of individual markers ensures that all protein positions are considered when accounting for compositional bias. Next, each of the filtered marker gene alignments were concatenated into a single MSA supermatrix for BMGE and one for Divvier, respectively, whereby previously removed gap-only sequences were added again in the corresponding positions. Finally, the MSA was trimmed according to GTDB criteria mentioned above to a length of 7859 amino acids (BMGE) and 32061 amino acids (Divvier) and used for phylogenetic inferences (see below).

### Phylogenetic inference

Phylogenomic trees were inferred with FastTree 2 (Price et al., 2010), ExaML (Kozlov et al., 2015), IQ-TREE (Nguyen et al., 2015), and PhyloBayes (Lartillot and Philippe, 2004) on different alignments and with a range of models (**Table S10**). Note that we chose IQ-TREE as the GTDB standard inference, since it scales with our dataset, allows mixture models (see below), and because a previous study concluded that for concatenation-based species tree inference, IQ-TREE consistently achieved the best-observed likelihoods for all data sets, compared to RAxML/ExaML and FastTree (Zhou et al., 2018). More details about each inference program is provided below.

#### FastTree

FastTree v2.1.9 was executed in multithreaded mode with the WAG+GAMMA parameters.

#### IQ-TREE

IQ-TREE was executed employing mixture models, such as C10-C60 (Le et al. 2008), and a faster approximation of these models known as “posterior mean site frequency” (PMSF) model (Wang et al., 2018). True C10-C60 trees are computationally more demanding, whereby memory requirements tend to increase with the number of components in the mixture, ranging from 10 to 60 (**Table S10**), and we therefore opted for the faster PMSF model, in particular C10 PMSF, to calculate the ar122.r89 tree. The tree was calculated with IQ-TREE v1.6.12 based on the C10 mixture model and a starting tree (-ft), inferred by FastTree v2.1.9 as described above, to invoke the faster PMSF approximation with the following settings: *-m LG+C10+F+G -ft <starting tree>*.

#### ExaML

ExaML trees (gamma, JTT) were calculated from 10 different starting trees with random seeds using the mpi version 3.0.20 (settings: -m GAMMA), whereby the tree with the highest likelihood score was retained.

#### PhyloBayes

To accommodate the computationally demanding Bayesian inferences we subsampled our data set by reducing the number of taxa to one representative per order resulting in 96 taxa. The order representatives were selected by removing genomes with a quality score (CheckM completeness - 5*CheckM contamination) of <50; <50% of the 122 archaeal marker genes; an N50 <4 Kb; >2,500 contigs or >1,500 scaffolds). From the remaining genomes, the highest quality genome was selected giving preference to i) NCBI reference genome annotated as “complete” at NCBI, ii) NCBI reference genomes, iii) complete NCBI representative genomes, iv) NCBI representative genomes, and v) GTDB representative genomes. **Table S19** indicates which of these categories a genome falls into. The Bayesian trees were inferred with PhyloBayes-MPI version 1.10.2 with the following settings: *-cat -gtr -x 10 -1 -dgam 4*. Three independent chains were run and tested for convergence (maxdiff =0.13) using bpcomp (part of pb_mpi) whereby the first 1,000 trees were eliminated as burn-in and the remaining trees were sampled every 10 trees.

#### Supertree

Supertrees were calculated from 1) 122 GTDB markers, and 2) from 253 markers which were present in >10% of archaeal genomes and were extracted from the PhyloPhlAn dataset (Segata et al., 2013). For the 122 marker set, a guide tree was inferred for each marker gene via approximate ML using FastTree v2.1.9 with the WAG model and gamma distributed rate heterogeneity. Subsequently, a complete ML tree was inferred using IQ-Tree v 1.6.9 with the C10 + PMSF model and the guide tree with 100 non-parametric bootstrap samplings to assess node support. A consensus tree was constructed from the individual gene trees using ASTRAL v5.6.3.

For the 253 marker set, we first identified, 400 conserved, single-copy gene orthologues from the PhyloPhlAn data set (Segata et al., 2013) using PhyloPhlAn v0.99. Markers present in less than 10% of all archaeal genomes were identified and removed resulting in a final marker set of 253 protein markers. For each marker, multiple-sequence alignment was performed using MAFFT-LINSI v7.221 (Katoh and Standley, 2013) and the resulting amino acid alignment was filtered using trimAl 1.2rev59, removing columns present in less than half of all taxa in the alignment. Tree inference was performed using the same approach as for the 122 marker set.

#### SSU trees

Trees from the trimmed SSU gene alignments (see section “SSU rRNA gene” above) were inferred with IQ-TREE, whereby the substitution model was determined by IQ-TREEs model finder to be SYM+R10.

### Taxonomic assignment and rank standardisation

The assignment of higher taxonomic ranks was normalized based on the relative evolutionary distance (RED) calculated from the ar122.r89 tree using PhyloRank (v0.0.37; https://github.com/dparks1134/PhyloRank/) as described previously (Parks et al., 2018). In brief, PhyloRank linearly interpolates the RED values of internal nodes according to lineage-specific rates of evolution under the constraints of the root being defined as zero and the RED of all present taxa being defined as one. To account for the influence of the root placement on RED values PhyloRank roots a tree multiple times, at the midpoint of each phylum with two or more classes. In the case of the ar122.r89 reference tree, PhyloRank identified 5 GTDB phyla (Halobacteriota, Thermoplasmatota, Methanobacteriota, Thermoproteota, and Asgardarchaeota) with two or more classes to be used for the multiple outgroup rooting approach. The RED of a taxon is then calculated as the median RED over all these tree rootings, excluding the tree in which the taxon was the outgroup. The RED intervals for each rank were defined as the median RED value ±0.1 to serve as a guide for the normalization of taxonomic ranks from genus to phylum in GTDB. The rank of species was assigned using ANI and AF criteria (see above). Note, that the application of names above the rank of genus in the GTDB taxonomy is manually curated where required, i.e. where naming ambiguity exists due to reclassification changes (taxa split, union, transfer) or where a new name needs to be assigned. Thereby, the curation team follows a decision tree for the manual curation workflow (**Fig. S21, Suppl. Text**).

### Assessment of phylogenetic congruence

We measured the phylogenetic tree similarity applying the normalized Robinson-Foulds distance (RF) of each alternative tree compared to the ar122.89 tree. The RF distance is defined as the number of splits that are present in one tree but not in the other one, and vice versa (Robinson and Foulds, 1981). The normalized RF is a relative measure obtained by dividing the calculated Robison-Foulds distance by the maximal RF. The resulting distance is a value between 0% and 100%, which can be interpreted as the percentage of different or missing splits in the alternative trees compared to the ar.122.89 tree (Kupczok et al., 2010). RF calculations and tree comparisons with lineage resolution were carried out with the visualisation tool metatree (https://github.com/aaronmussig/metatree).

### Assessment of taxonomic congruence

The congruence of the GTDB taxonomy in different trees was assessed as (i) the percentage of taxa identified as monophyletic, operationally monophyletic, defined as having an F measure ≥0.95, or polyphyletic, (ii) the RED distributions for taxa at each rank relative to the median RED value of that rank, and (iii) the number of genomes with identical or conflicting taxonomic assignments between compared trees. Thereby (i) was carried out by placing each taxon on the node with the highest resulting F measure, which is defined as the harmonic mean of precision and recall, and it has been proposed for decorating trees with a donor taxonomy (McDonald et al., 2012). Note, we introduced the term operationally monophyletic (F measure ≥0.95), since otherwise a few incongruent genomes can cause a large number of polyphyletic taxa.

## Data availability

The GTDB taxonomy is available at the GTDB website (https://gtdb.ecogenomic.org/), including the ar122.r89 tree and the GTDB and NCBI taxonomic assignments for all 2,392 archaeal genomes in GTDB 04-RS89. The standalone tool GTDB-Tk, which enables researchers to classify their own genomes according to the GTDB taxonomy is available at GitHub (https://github.com/Ecogenomics/GTDBTk/) and through KBase (https://kbase.us/applist/apps/kb_gtdbtk/run_kb_gtdbtk/release). Genome assemblies are available from the NCBI Assembly database (BioProject: PRJNA593905).

## Acknowledgements

We thank Brian Kemish and Dinindu Senanayake for system administration support, Pelin Yilmaz for stimulating discussions on archaeal taxonomy, and the GTDB user community for their feedback. We also thank the Australian Centre for Ecogenomics (ACE) at The University of Queensland and the New Zealand eScience Infrastructure (NeSI) for providing high performance computing facilities. The project was supported by an Australian Research Council (ARC) Future Fellowship (FT170100213) awarded to C.R. and by an Australian Research Council Laureate Fellowship (FL150100038) awarded to P.H.

## Supplementary information

### Supplementary Tables and Figures

See files “*Supplementary Tables” 1 and 2, and “Supplementary Figures*”.

### Supplementary data

All GTDB files for Release 04-RS89 are available through the GTDB repository https://data.ace.uq.edu.au/public/gtdb/data/releases/release89/89.0/

### Supplementary File 01 | Phylogenetic trees

Newick files of all GTDB decorated trees: *Suppl_file_01_trees_4_paper*.*zip*

### Supplementary File 02 | SR4 model used for data recoded inferences

File name: *\**.*nex*

## Notes

https://data.ace.uq.edu.au/public/gtdb/data/releases/release89/89.0/

### Competing Interest Statement

The authors have declared no competing interest.

